# Neuropathological and Functional Impact of Astrocyte-Derived Extracellular Vesicles in an Aged Model of Alzheimer’s Disease

**DOI:** 10.64898/2026.02.13.705826

**Authors:** Bao Quach, Sahar Salehi, Rejina Roufegarinejad, Michael Mante, Jazmin Florio, Robert A. Rissman, Charisse N. Winston

## Abstract

Extracellular vesicles (EVs) are lipid-bound particles that transfer cargos between cells. While plasma neuronal-derived EVs (NEVs) from individuals with mild cognitive impairment (MCI) and Alzheimer’s disease (AD) have been reported to exhibit high pathogenic potential, this study examined the impact of astrocyte-derived EVs (AEVs) in an aged AD mouse model. Plasma AEVs were isolated from cognitively normal control (CNC), MCI, and AD individuals using GLAST-based immunocapture and AEV size, purity, and tetraspanin were validated by flow cytometry, nanoparticle tracking, and super-resolution microscopy. AEVs pooled by clinical cohort were injected into the hippocampus of 6-month-old female PSAPP mice. Behavioral, biochemical, and neuropathological outcomes were assessed 6 months later. Rotarod assessment revealed significant impairment in motor coordination (*p*<0.0001) in mice receiving MCI- and AD-AEVs compared with those receiving CNC-AEVs. Morris water maze (MWM) also demonstrated cognitive deficits in MCI- and AD-AEV injected mice; however, no overt changes were observed in the staining of amyloid plaque burden (6E10), and neuroinflammation (GFAP). Immunoblotting of 82E1 and 22C11 confirmed Aβ/AAP levels remained similar across all injected mice, whereas increased cortical tau accumulation was observed in MCI- and AD-AEV injected mice. Cerebellar synaptic density (SY-38) remained unchanged. While plasma AEVs from healthy individuals may confer neuroprotective benefits, AEVs from cognitively impaired individuals promoted tau accumulation in an amyloidogenic mouse model. These findings suggest that AEVs may play a dual role in AD as both potential biomarkers and mechanistic drivers of neurodegeneration, highlighting their relevance as targets for future therapeutic strategies.

## Introduction

In Alzheimer’s disease (AD), Aβ is primarily produced through proteolytic cleavage of amyloid precursor protein (APP) in neurons, while astrocytes have been reported to contribute to a relatively lesser extent (Hampel et al., 2021). The pathogenicity of astrocytes and their involvement in Aβ production are not yet fully understood. Astrocytes are a major subset of glial cells that are vital for maintaining the central nervous system (CNS). In a healthy CNS, astrocytes perform critical neuronal functions, including regulating ion homeostasis (Kim et al., 2019), interacting with endothelial cells that comprise the blood-brain barrier (BBB) (Abbott et al., 2006), and facilitating synapse formation by eliminating synaptic debris (Kim et al., 2019). While astrocytes maintain CNS architecture under healthy conditions, substantial evidence indicates that they can also facilitate neurotoxicity and contribute to pathogenesis in neurodegenerative disorders such as AD (Phatnani and Maniatis, 2015; Zhao et al., 2021). In AD, astrocytes have been shown to participate in Aβ clearance (Liu et al., 2017). However, reactive astrocytes can propagate feed-forward signals that interact with aging and degenerating neurons, microglia, and other CNS cells, ultimately promoting neurodegeneration (Preman et al., 2021). Therefore, the identification of elevated astrocytic biomarkers that predict conversion from mild cognitive impairment (MCI) to AD is critical for improving early diagnosis.

Targeting the extracellular vesicles (EVs) released by astrocytes represents a viable approach to mitigating astrocyte-driven pathogenesis. EVs are cell-derived, lipid bilayer-bound vesicles that mediate intercellular communication in the CNS by transporting proteins, lipids, and nucleic acids. EVs are typically categorized into three different subtypes: exosomes (40–150 nm), microvesicles (50–2,000 nm), and apoptotic bodies (50–5,000 nm) (Yáñez-Mó et al., 2015). Under homeostatic conditions, EVs contribute to CNS maintenance by delivering neuroprotective, antioxidant, and anti-inflammatory molecules that modulate neuroinflammatory response. They also facilitate synaptic plasticity and participate in the clearance of metabolic byproducts and aggregated proteins (Phatnani and Maniatis, 2015; Zhao et al., 2021).

Recent studies in AD have reported that EV functions in the CNS are context dependent. EVs are released by nearly all cell types, and their cargo can be indicative of cellular origin and physiological state. Under pathological conditions, changes in rate and content of EVs release have been observed in multiple CNS cell types with evidence that inflammatory and stress signals can modulate their secretion and cargo (Khan et al., 2022; Thompson et al., 2016). EVs have also been researched extensively in regard to their contribution to CNS disorders and have been identified as biomarkers in CNS-related injuries (Zhao et al., 2021).

Previous studies demonstrate that plasma neuronal-derived EVs (NEVs) from individuals with MCI and AD can induce tau accumulation and neurodegeneration in the brains of recipient mice, indicating the pathogenic potential of circulating NEVs to facilitate the spread of toxic tau proteins (Winston et al., 2019a; Winston et al., 2016). Additionally, Aβ released by microglial-derived EVs (MEVs) have been reported to contribute to early synaptic dysfunction in vitro (Gabrielli et al., 2022). Building on these observations, we reasoned that other EV populations, specifically astrocyte-derived EVs (AEVs), may carry cargo capable of influencing disease progression. AEVs from AD patients are known to contain defined molecular cargo, including complement components and other regulators of neuroinflammation (Upadhya et al., 2020; Winston et al., 2019b). Moreover, they are implicated in tauopathies through their ability to carry and accumulate tau isoforms. However, their specific role in tau propagation remains unclear, highlighting the need to evaluate AEV effects in vivo (Perbet et al., 2023). Recent studies have also shown astrocytes and AEVs implicated in TDP-43 proteinopathies in dementia-related diseases such as amyotrophic lateral sclerosis (ALS), frontotemporal dementia (FTD), and limbic predominant age-related TDP-43 encephalopathy (LATE), further underscoring the role of astrocytes and AEVs in pathological signaling ((Winston et al., 2022); Dellar *et al*., 2025; Velebit *et al*., 2020; Ng and Ng, 2022).

We therefore investigated how human plasma AEVs are delivered and trafficked in the brain and assessed their neuropathological and functional impact in an amyloidogenic mouse model. We employed the APPswe x PSEN1dE9 (PSAPP) mice, which have been shown to develop early and robust Aβ plaque deposition and age-dependent cognitive deficits (Borchelt et al., 1997; Kim et al., 2015; Zhong et al., 2024). Notably, the model exhibits AD-relevant behavioral deficits in an age-dependent manner. This mouse model develops measurable impairments in memory as early as 6 months of age, with deficits in spatial navigation and learning by 12 months of age (Janus et al., 2015; Lalonde et al., 2005; Prikhodko et al., 2020). However, PSAPP mouse models also do not recapitulate the full spectrum of human AD, particularly neurofibrillary tangles (NFTs) and overt tauopathy, limiting direct translational inference (Yokoyama et al., 2022; Zhong et al., 2024). Because PSAPP mice are primarily amyloidogenic and do not spontaneously develop robust tau pathology, they provide a ‘blank slate’ platform to isolate the effects of AEVs. This background is advantageous for investigating whether human AEVs can precipitate the earliest tau phosphorylation events observed in prodromal AD, rather than merely amplifying existing tangles (Jankowsky and Zheng, 2017; Kitazawa et al., 2012). We hypothesized that, analogous to NEVs, AEVs from individuals with MCI or AD would exhibit higher pathogenic potential than AEVs from CNC individuals. To test this, we administered AEVs to PSAPP mice at 6 months of age, where amyloid pathology emerges, and subsequently evaluated behavioral and neuropathological outcomes.

## Materials and Methods

### Patient Demographics

Plasma samples were acquired through the University of California, San Diego (UCSD) Shiley-Marcos Alzheimer’s Disease Research Center (ADRC) (Winston et al., 2016). Pooled EV preparations were generated from three clinical cohorts: cognitively normal controls (CNC, n = 4; mean age, 77.5 (11.22); patients with stable diagnosis of mild cognitive impairment (MCI; n = 4, mean age 69.5 (14.25) and patients with an established diagnosis of mild to moderate AD (AD, n = 4, mean age 70.25 (9.12), and CNC, MCI, and AD groups were initially defined based on CSF Aβ concentrations and cognitive status as indicated by a mini-mental state examination score (MMSE): CNC, CSF Aβ1-42 level >190 pg/mL, and normal cognition (MMSE 27–30); MCI, memory complaint, cutoff score on the Logical Memory test, MMSE >24, intact ADL; AD, CSF Aβ1-42 <190 pg/mL and MMSE <20 (Grundman et al., 2004).

### Isolation and Enrichment of GLAST-Positive Astrocytes Extracellular Vesicles (AEVs) From Human Plasma via Fluorescence Activated Cell Sorting (FACS) and Bead Antibody Exosome (BAE)-FITC Complex

EVs isolation was performed as described in the manufacturer’s protocol (System Biosciences, Inc., Mountain view, CA, United States; Catalog # EXOQ5TM-1). First, 2.5 µL purified thrombin (System Biosciences, Inc.; Catalog # TMEXO-1) was added to 250 µL of human plasma and incubated for 5 minutes at room temperature. The resultant mixture was centrifuged at 10,000 rpm for 5 minutes. 63 µL ExoQuick Exosome Precipitation solution (System Biosciences, Inc.; Catalog # EXOQ5TM-1) was added to the supernatant, and the mixture was incubated for 1 hour at 4℃ and centrifuged for 1 hour at 1500 x g at 4℃. The resultant pellet was resuspended in 300 µL of 1X phosphate buffer saline (PBS) (Thermo Fisher Scientific; Catalog # AM9625) with Halt protease and phosphatase inhibitor cocktail EDTA-free (Thermo Fisher Scientific; Catalog # 78443). Samples were stored at -80℃ until enrichment of EVs from astrocytes. Particle identity was confirmed by nano-flow cytometry (NanoFCM, NanoFCM Inc.). Samples were accepted only if they showed a modal diameter of 100–150 nm, GLAST+ events > 80 %, calnexin+ events < 1 %, and a particle/protein ratio > 3 × 109 particles µg-1, mirroring criteria established in our prior TEM-validated studies (Goetzl et al., 2016;(Winston et al., 2019c);(Winston et al., 2022); (Figueroa-Hall et al., 2025).

Enrichment of AEVs was performed as described in the manufacturer’s protocol using 9.1 um streptavidin magnetic Exo-Flow beads (System Biosciences, Inc.; Catalog # CSFLOWBASICA-1). To start, 45 µL of these beads were incubated with mouse anti-human GLAST (ACSA-1) biotinylated antibody (Miltenyi Biotec, Inc., Auburn, CA, United States; Catalog # 130-118-984) for 2 hours on ice. The bead-antibody (bead-Ab) complexes were gently flicked every 30 minutes to ensure mixing. 1X Bead Wash Buffer (System Biosciences, Inc.; CSFLOWBASICA-1) was added to the bead-Ab complexes and washed three times using a magnetic stand. The bead-Ab complexes were then suspended in 300 µL of 1X Bead Wash Buffer and 100 µL of total EVs suspension, rotating overnight at 4℃. After, the bead-Ab-EVs (BAE) complexes were washed three times with 1X Bead Wash Buffer and then suspended in 240 µL of Exosome Stain Buffer and 10 µL of Exo-FITC Exosome FACS stain (Systems Biosciences, Inc.; Catalog # CSFLOWBASICA-1) for 2 hours on ice. The BAE-FITC complexes were gently flicked every 30 minutes to ensure mixing. The complexes were subsequently washed three times in 1X Bead Wash Buffer and then suspended in 300 µL of 1X Bead Wash Buffer prior to loading into BD FACS Aria II for sorting.

The BAE-FITC complexes were incubated with 300 µL of Exosome Elution Buffer (System Biosciences, Inc.; Catalog # FLOWBASICA-1) at 25℃ for 30 minutes. Lastly, the supernatant with the eluted EVs were incubated with 1 µL of Exo-FlowIP clearing reagent (System Biosciences, Inc.; Catalog # EXOFLOW32A) at 37℃ for 30 minutes, then stored at -80℃. Nanoparticle tracking analysis (NTA) was used to validate plasma AEVs by confirming size and concentration. Super-resolution imaging was performed with the ONI Nanoimager “EV Profiler” kit (CD9/63/81). GLAST-specific probes are not yet commercially available; therefore astrocytic origin was confirmed by anti-GLAST magnetic enrichment followed by western blot and FACS (see Fig. 2A–C). Preparations were accepted only when GLAST⁺ events ≥ 80 % and calnexin/GM130⁺ events ≤ 1 %. Purity safeguards included: (i) ExoQuick Ultra resin cleanup to deplete soluble plasma proteins; (ii) anti-GLAST bead capture followed by three 10 mL washes in 0.1 % Tween-20; (iii) acceptance threshold of ≥ 1 × 10¹L particles µg⁻¹ protein (NTA/BCA); and (iv) western blot and FACS panels showing strong CD9/63/81 and GLAST with negligible calnexin, GM130, or ApoA1. To obtain the ≥ 10 µg protein required for each intracerebral injection, plasma from four age-, sex-, and APOE-matched donors was pooled per clinical group. Single-donor yields are ≤ 0.3 µg per 250 µL after GLAST immunocapture(Figueroa-Hall et al., 2025), making individual dosing impracticable under IRB-approved blood-draw limits.

**Figure 1:**
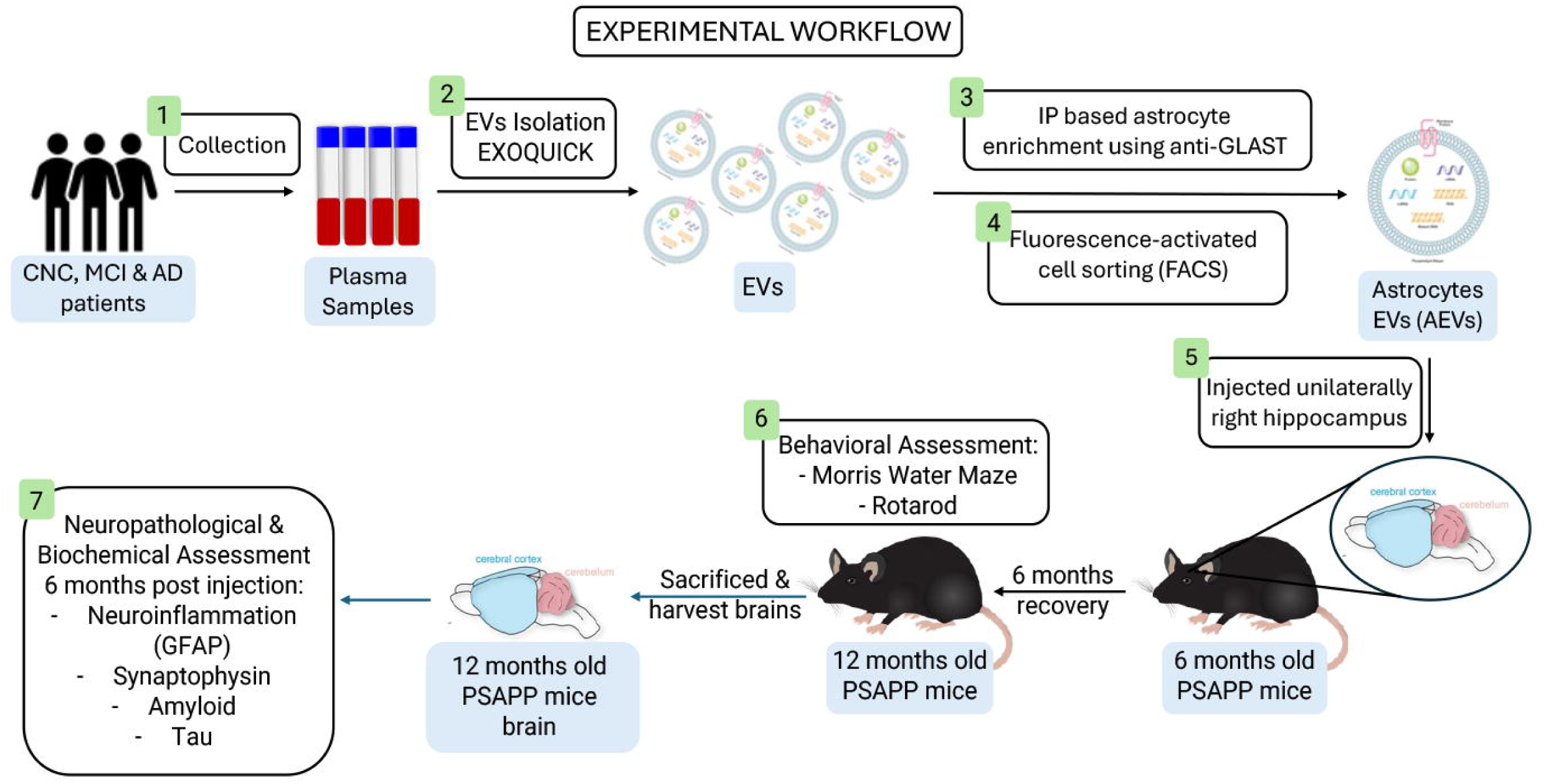
A representative experimental workflow describing the steps of the study. Extracellular vesicles (EVs) were isolated from human plasma of cognitively normal control (CNC), mild cognitive impaired (MCI), and Alzheimer’s disease (AD) individuals and astrocyte-derived EVs (AEVs) were enriched using anti-human GLAST immunocapture. AEVs were pooled by clinical cohort and unilaterally injected into the right hippocampus of 6-month-old female PSAPP mice with a 6-month convalescent period. Behavioral assessments were done at 6-months post injection (mpi) (Morris Water Maze & Rotarod). Mice were then sacrificed, and brains were harvested for neuropathological and biochemical assessment (amyloid, tau, synaptic density, and neuroinflammation). Abbreviations: CNC, cognitively normal control; MCI, mild cognitive impairment; AD, Alzheimer’s disease; EVs; extracellular vesicles; IP, immunoprecipitation; PSAPP, APPswe x PSEN1dE9.

**Figure 2:**
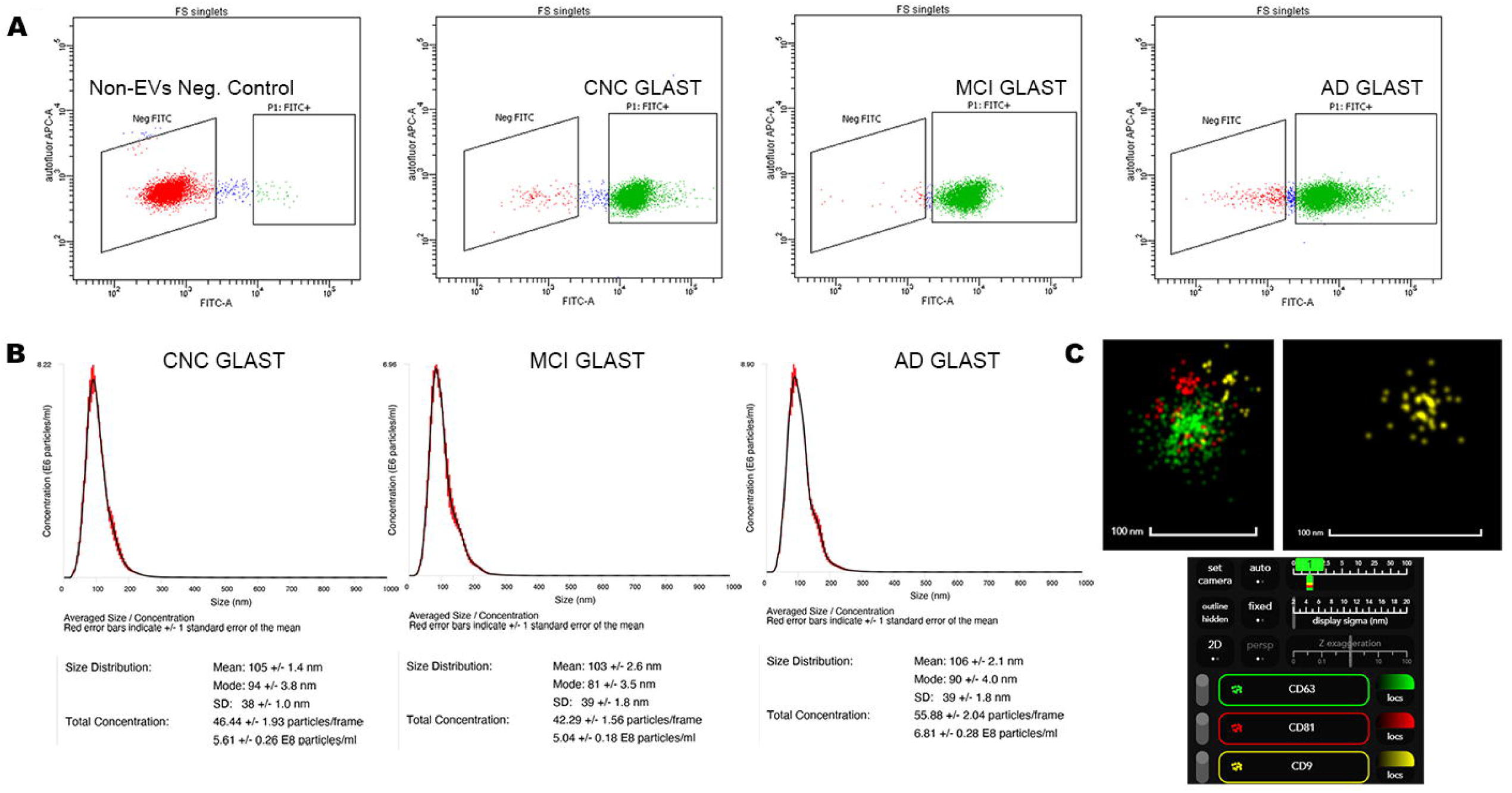
Isolation and enrichment of GLAST-positive astrocyte-derived extracellular vesicles (AEVs) from human plasma via fluorescence activated cell sorting (FACS) and validated with nanoparticle tracking analysis (NTA) and super resolution microscopy (ONI Nanoimager). (A) Representative FACS plot for non-exosomes, negative control (red) and bead-antibody-exosome (BAE)-FITC complexes generated from EVs (green) isolated from a cognitively impaired individual and enriched against anti-GLAST antibody. (B) Representative plot of size and concentration determined by NTA for isolated plasma EVs. (C) Representative image of plasma-derived EV positive for extracellular markers: CD63, CD81, and CD9 determined by super-resolution microscopy (ONI Nanoimager). Representative image of a single plasma-derived AEV positive for CD9. Abbreviations: EVs, extracellular vesicles; CNC, cognitively normal control; MCI, mild cognitive impairment; AD, Alzheimer’s disease.

**Table 1.** Patient demographics showing age, sex, APOE4 status, and race. Abbreviations: CNC, cognitively normal control; MCI, mild cognitive impairment; AD, Alzheimer’s disease.

|  | CNC | MCI | AD |
| --- | --- | --- | --- |
| Age |  |  |  |
| Mean (SD) | 77.5 (11.22) | 69.5 (14.25) | 70.25 (9.12) |

| **Sex** |  |  |  |
| --- | --- | --- | --- |
| Male | 2 (50%) | 3 (75%) | 2 (50%) |
| Female | 2 (50%) | 1 (25%) | 2 (50%) |
| **APOE4** |  |  |  |
| Non-Carrier | 2 (50%) | 2 (50%) | 1 (25%) |
| Carrier | 2 (50%) | 2 (50%) | 3 (75%) |
| **Race** |  |  |  |
| White | 4 (100%) | 4 (100%) | 4 (100%) |
| Black | 0 (0%) | 0 (0%) | 0 (0%) |
| Hispanic | 0 (0%) | 0 (0%) | 0 (0%) |

### Unilateral Injection of GLAST-Positive AEVs From Plasma Into PSAPP Mice Hippocampus

PSAPP mice (B6C3-Tg (APPswe, PSEN1dE9) 85Dbo/Mmjax) co-express mutant human APP (Swedish, K670N/M671L) and PSEN1 (ΔE9). They begin forming Aβ plaques deposition beginning at ∼6 months of age, with progressive amyloid accumulation by 9–12 months (Garcia-Alloza et al., 2006; Jankowsky et al., 2004) (JAX #004462). We therefore used aged cohorts (9–12 months), a stage that combines robust Aβ pathology with minimal endogenous tauopathy, creating a sensitive background for detecting AEV-induced tau phosphorylation or aggregation. At these ages the mice also exhibit AD-relevant behavioral deficits, including age-dependent impairments in spatial learning and memory tasks. Mice were housed under standard conditions. Procedures were performed under the approval of the IACUC of the University of California, San Diego (UCSD) under the protocol #S02221. All efforts were made to minimize the suffering of the animal and to reduce the number of animals used.

The following setup for mouse intrahippocampal injection was performed as described (Taylor et al., 2021), with a few modifications. The vaporizer was set to 3% isoflurane rather than 2%. Shortly after anesthetization, buprenorphine (0.05 mg/kg) was administered subcutaneously for analgesia. Following fur removal, the surgical area was sterilized with povidone iodine. A 10–15 mm incision was made to expose the skull, and the area was dried prior to injection. A 10 µL syringe (Hamilton; Catalog # 80366 26s gauge) was mounted to the apparatus and positioned at the bregma, with the coordinates zeroed using the digital display. The following coordinates used for the right hippocampal injection were: -2.0 mm in the anteroposterior axis, +1.5 mm in the mediolateral axis, and -1.3 mm in the dorsoventral axis. The desired location was marked and subsequently drilled. The dorsoventral axis was set at 0.0 with the needle tip aligned over the drill hole. It was then lowered slowly to -3.0 mm in the dorsoventral axis. The infusion pump was then set to administer 2 µL of prepared AEVs (2.5 ug/uL) at a rate of 1 µL/min. Once the infusion was complete, the needle remained in place for an additional minute before being withdrawn. The incision was sealed with tissue adhesive (GLUture), and the mouse was removed from the apparatus and returned into its respective cage. Mice were monitored throughout the 6 months post-injection period.

### Behavioral Assessment

#### Morris Water Maze (MWM)

Cognitive performance was assessed using the Morris water maze (MWM) as previously described (Ngolab et al., 2021). Testing spanned eight consecutive days (Monday–Friday, then Monday–Wednesday) with 24-hour intersession intervals (weekends excluded). A circular pool (diameter, 180 cm) was filled with water maintained at 24 °C and rendered opaque with nontoxic white paint. The pool was divided into four quadrants; the escape platform was fixed in the northwest (NW) quadrant for the duration of testing. During cued (visible platform) trials, the platform was marked with a pole wrapped in alternating black-and-white tape.

From Days 1–7, mice completed four trials per day (maximum trial duration, 90 seconds; inter-trial interval, 2–3 minutes). On Day 1, all trials began from the west start point. Mice that failed to locate the platform on their first trial were gently guided to it and allowed to remain for 30 seconds; this assistance was not repeated on subsequent trials. On Days 2–3, start positions alternate between the south and east points. Cued training occurred on Days 1–3. On Days 4–7 (hidden platform phase), the cue was removed, and the platform was submerged to assess spatial learning.

On Day 8, a two-trial probe was conducted. In Trial 1 (platform removed; start = south), mice swam for 40 seconds to assess memory retention. Primary probe metrics included entries (crossings) into the former platform location; we also quantified time spent in each quadrant, entries into the target quadrant, and total swim distance. In Trial 2 (platform and cue restored; start = east), mice were given 40 seconds to locate the platform.

### Rotarod

Motor coordination and balance were assessed on an accelerating five-lane rotarod (Ugo Basile, model 47650) over two days (Podvin et al., 2025). On Day 1, mice completed two acclimation trials (Trial 1: acceleration from 5–10 rpm over 120 seconds; Trial 2: from 5–20 rpm over 240 seconds; 10-minute inter-trial interval), followed by three performance trials (Trials 3–5: acceleration from 5–40 rpm; maximum duration 240 seconds; 10-minute inter-trial interval). On Day 2, mice completed seven additional performance trials (acceleration from 5–40 rpm; maximum duration 240 seconds; 10-minute inter-trial interval). The apparatus automatically recorded latency to fall (seconds) and speed at fall (rpm). If a mouse remained on the rod for the full trial, the maximum trial time (120 seconds for Day 1, Trial 1; 240 seconds otherwise) was assigned. Brief passive rotations (i.e., clinging for one full revolution without active stepping) were scored as a fall. On the test day, mice were transported to the behavior room in their home cages, weighed and ID-marked (tail color-coded), and the rod surface and lanes were cleaned with 70% isopropanol between mice.

### Brain Harvest, Immunochemistry and Immunoblot

Mice were euthanized 24 hours after the final behavioral assessment by transcardial perfusion with saline in accordance with NIH and IACUC guidelines. Brains were either snap-frozen in liquid nitrogen for biochemical analyses or immersion-fixed in 4% phosphate-buffered formaldehyde (PFA) for 48 hours at 4 °C. Fixed tissue was subsequently cryoprotected in 20% sucrose in 1X phosphate-buffered saline (PBS) for 24 hours at 4 °C. For neuropathological assessment, frozen brains were coronally sectioned at 40 µm using a freezing-sliding microtome. Free-floating sections were incubated overnight at 4 °C with primary antibodies: mouse anti-6E10 (BioLegend, Cat# 803003), mouse anti-GFAP-biotin conjugate (Millipore, Cat# MAB3402B), or mouse anti-phospho-tau (Ser202/Thr205, AT8; Invitrogen, Cat# MN1020). Sections were then incubated with biotinylated horse anti-mouse IgG secondary antibody (Vector Laboratories, Cat# BA-2000), except for GFAP which was already biotin-conjugated, followed by detection using the avidin-biotin complex method (VECTASTAIN Elite ABC Kit, Peroxidase Standard, Vector Laboratories, Cat# PK-6100). Immunoreactivity was visualized with diaminobenzidine (DAB) chromogen (Vector Laboratories, DAB Substrate Kit, Cat# SK-4100). Slides were imaged using a NanoZoomer S60 Digital Slide Scanner (Hamamatsu) at 20X magnification.

For synaptic density assessment, free-floating sections were incubated overnight at 4 °C with mouse anti-synaptophysin (SY-38; Millipore, Cat# MAB5258-I). Sections were then incubated with biotinylated horse anti-mouse IgG followed by streptavidin-HRP and subsequently labeled with tyramide red diluted in amplification diluent (Akoya Biosciences, Cat# NEL702001KT). Nuclei were counterstained with Hoechst (Invitrogen, Cat# 447167), and sections were mounted and cover-slipped using ProLong Gold Antifade Mountant with DAPI (Invitrogen, Cat# P36931). Imaging was performed using a Leica Stellaris 8 confocal microscope.

For biochemical analysis, frozen brains from three mice per group were dissected into the cortex and hippocampus. Tissue was homogenized in 1X nuclease-free PBS containing protease inhibitors (Mini-Complete, Roche) using a Bead Mill 24 (Fisher) with 1.4 mm ceramic beads (Fisher). Homogenates were incubated in 10X RIPA buffer, and protein concentrations were quantified using a BCA assay (Bio-Rad). Equal amounts of protein (20 µg per lane) were separated on 4–20% gradient SDS-PAGE gels (Criterion, Cat# 5678094) and transferred onto PVDF membranes (Millipore, Cat# ISEQ00010) using the semi-dry Trans-Blot Turbo Transfer System (Bio-Rad). Membranes were probed with the following primary antibodies: mouse anti-APP (22C11; Millipore, Cat# MAB348), mouse anti-Aβ (6E10; BioLegend, Cat# 803003), mouse anti-APP C-terminal (82E11; IBL-America, Cat# 10323), mouse anti-phospho-tau (PHF-1; gift from Dr. Peter Davies), and rabbit anti-β-actin (Cell Signaling Technology, Cat# 5125S). Blots were incubated with species-appropriate HRP-conjugated secondary antibodies (Bio-Rad, Cat# 172-1019 and Cat# 172-1011), and signals were visualized by enhanced chemiluminescence (SuperSignal West Pico PLUS, Thermo Scientific).

### Immunohistochemical Quantification

Quantification of 6E10 and GFAP brightfield staining intensity was performed using FIJI (ImageJ). To minimize variability between staining batches, the corpus callosum, which showed consistent signal across samples, was used as a reference region. The optical density (OD) of the corpus callosum, representing the lightest background signal, was subtracted from the OD of the cortex and hippocampus to generate standardized values for these regions of interest (ROIs). Standardized OD values were graphed and statistically analyzed for significance.

For fluorescence labeling, FIJI was also used to quantify DAPI and synaptophysin signal intensity. Within each ROI, the mean OD was calculated from three sample fields. To normalize synaptophysin signal, the mean OD of synaptophysin was divided by the corresponding mean OD of DAPI.

### Immunoblot Quantification

Immunoblot images were analyzed using Image Lab Standard Edition software (Bio-Rad). Protein bands were selected within each lane and manually resized to correspond precisely to the area of each detected band. Bands at the expected molecular weight (kDa) of interest were identified, and lane profile plots were used to quantify band intensity and total protein signal. The software used generated an analysis table in which the intensity of each band was calculated in arbitrary units, with adjusted volume representing the relative protein amount. Normalization was performed in two steps. First, the adjusted volume of each band was normalized to the β-actin loading control by dividing each adjusted value by the highest β-actin signal. Second, for each antibody of interest, the adjusted band intensity was divided by this normalization factor to yield a normalized signal per lane. For all antibodies except β-actin (control), the mean normalized signal was calculated across the CNC-, MCI-, and AD-AEV injected groups for statistical comparison.

### Statistical Analyses

All statistics were performed in GraphPad Prism v10.0 (GraphPad Software, San Diego, CA, USA). Data distributions were first examined with the Shapiro–Wilk test to guide the choice of parametric versus non-parametric procedures. Group differences involving two independent samples were evaluated with an unpaired Student’s t-test when normality held or a Mann–Whitney U test when it did not. Experiments with three or more independent groups were analyzed by one-way ANOVA; if the overall F value reached significance, pairwise contrasts were resolved with Holm–Šídák post-hoc testing to control the family-wise error rate. Repeated-measures designs (e.g., longitudinal behavioral readouts) were assessed with two-way repeated-measures ANOVA. Results are expressed as mean ± SEM, and figure legends include exact p values and effect sizes (Cohen’s d for two-group tests or η² for ANOVA). Statistical significance was defined as two-tailed p < 0.05.

## Results

### Characterization of Plasma of AEVs using FACS, Nanoparticle Tracking Analysis (NTA), and super-resolution imaging

To confirm successful isolation and enrichment of AEVs from human plasma, final bead-Ab-EVs (BAE) complexes prepared from all mice groups were tagged with a FITC-conjugated antibody (Exo-FITC, System Biosciences Inc.) and analyzed by fluorescence activated cell sorting (FACS). Bead-antibody-FITC (BA-FITC) complexes, which lack EVs, served as negative controls to verify the absence of non-specific Exo-FITC binding.

Fluorescent activated cell sorting FACS analysis demonstrated that AEVs prepared from CNC, MCI, and AD human plasma were 65.3%, 61.7%, 59.4% Exo-FITC positive, respectively (n = 3). In contrast, less than (0.2%) of the Exo-FITC positive signal was captured in the non-EVs, negative controls (BA-FITC) (Figure 2A). NTA further validated the size distribution and concentration of the enriched AEVs, revealing the particle size to be between 80–100 nm (Figure 2B). Super-resolution imaging with the ONI Nanoimager confirmed the presence of canonical EV markers: CD63, CD81, and CD9 (Figure 2C).

### Assessment of the Learning and Memory in AEV-injected PSAPP mice

Spatial memory was evaluated at 6 mpi in all AEV-injected PSAPP mice using the MWM, as previously described (Figure 3). During cued (visible platform) training (Days 1–3), MCI- and AD-AEV injected mice showed longer escape latencies and greater path lengths than the CNC-AEV injected mice. Across the hidden (submerged) platform phase (Days 4–7), performance of all three groups converged; by Day 7, groups exhibited similar escape latencies and path lengths to reach the platform (Figure 3A–B). In the probe trial (Day 8, Trial 1; platform removed, 40 seconds), MCI- and AD-AEV injected mice made fewer entries (crossings) into the former platform location compared with CNC-AEV injected mice (Figure 3C & 3E). In Trial 2 (platform restored; 40 seconds), all groups showed comparable ability to locate the platform, with similar escape latencies (Figure 3D) and path lengths (Figure 3F).

**Figure 3:**
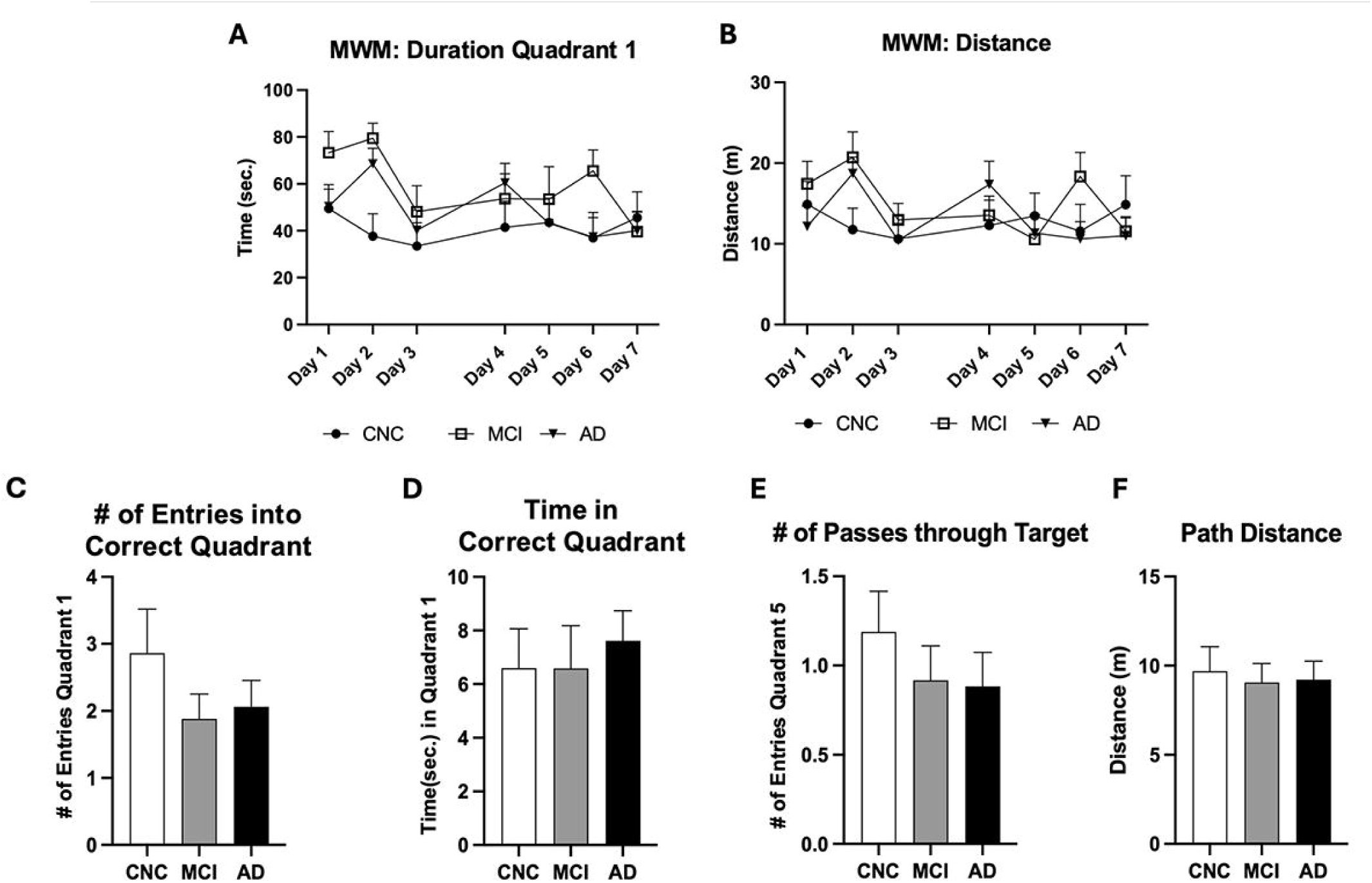
Morris Water Maze (MWM) behavioral assessment of astrocyte-derived extracellular vesicles (AEVs) unilaterally injected into the right hippocampus of female PSAPP mice. (A) Displays the duration (seconds) in quadrant 1 and (B) the distance (meters) travelled across the seven-day trial. (C) The number of entries into the correct quadrant, (D) the amount of time (seconds) spent in the correct quadrant, (E) the number of passes through the target (platform), and (F) the path distance (meters) travelled on Day 8, the probe trial. (n = 10–14/group, female, 6 months post-injection (mpi)). Statistical analyses were performed using One-way ANOVA with a Tukey test. All comparisons were not significant. Abbreviations: CNC, cognitively normal control; MCI, mild cognitive impairment; AD, Alzheimer’s disease; PSAPP, APPswe x PSEN1dE9.

### Histological Assessment of Amyloid burden and Inflammation in AEV-injected PSAPP Mice

Aggregated Aβ plaques (Figure 4) and inflammation (Figure 5) were quantified using 6E10 and GFAP immunostaining, respectively, in the ipsilateral and contralateral hemispheres of AEV injected PSAPP mice. CNC-AEV injected PSAPP mice showed lower cortical and hippocampal aggregated Aβ plaques compared to MCI- and AD-AEV injected PSAPP mice. Notably, MCI-AEV injected mice exhibited higher plaque expression on the ipsilateral side compared to the AD-AEV injected mice (Figure 4C–D). However, these differences did not reach statistical significance. Cortical astrocyte reactivity and neuroinflammation was reduced in CNC-AEV injected mice compared with MCI-AEV injected mice, whereas AD-AEV injected mice showed levels comparable to CNC-AEV injected mice, although these differences were not statistically significant (Figure 5C). In the hippocampus, CNC-AEV injected mice exhibited greater astrocyte reactivity and neuroinflammation on the ipsilateral side than the AD-AEV injected mice, while the MCI-AEV injected mice exhibited the highest levels, though again, no significant differences were observed. On the contralateral side, astrocyte activity and neuroinflammation were similar across all three AEV-injected mouse groups (Figure 5D).

**Figure 4:**
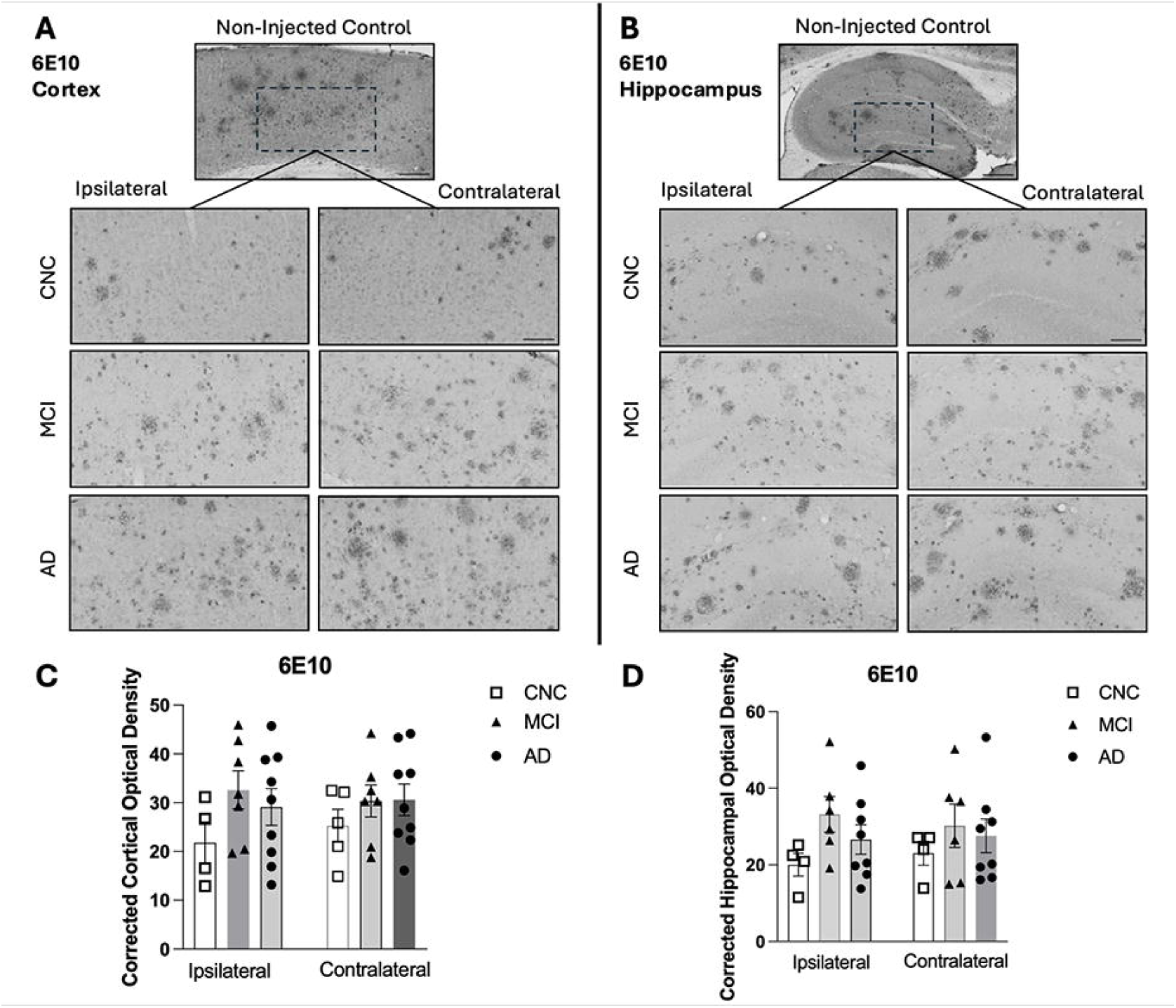
6E10 brightfield immunohistochemical staining of astrocyte-derived extracellular vesicles (AEVs) unilaterally injected into the right hippocampus of female PSAPP mice at 6 months post-injection (mpi). Representative staining of 6E10 in the (A) cortex and (B) hippocampus. Scale bars = 250 μm, 150 μm. Depicts the corrected optical densities of 6E10 staining in the (C) cortex and (D) hippocampus. At least four mice per group were analyzed. Statistical analyses were performed using One-way ANOVA with a Tukey test. All comparisons were not significant. Abbreviations: CNC, cognitively normal control; MCI, mild cognitive impairment; AD, Alzheimer’s disease.

**Figure 5:**
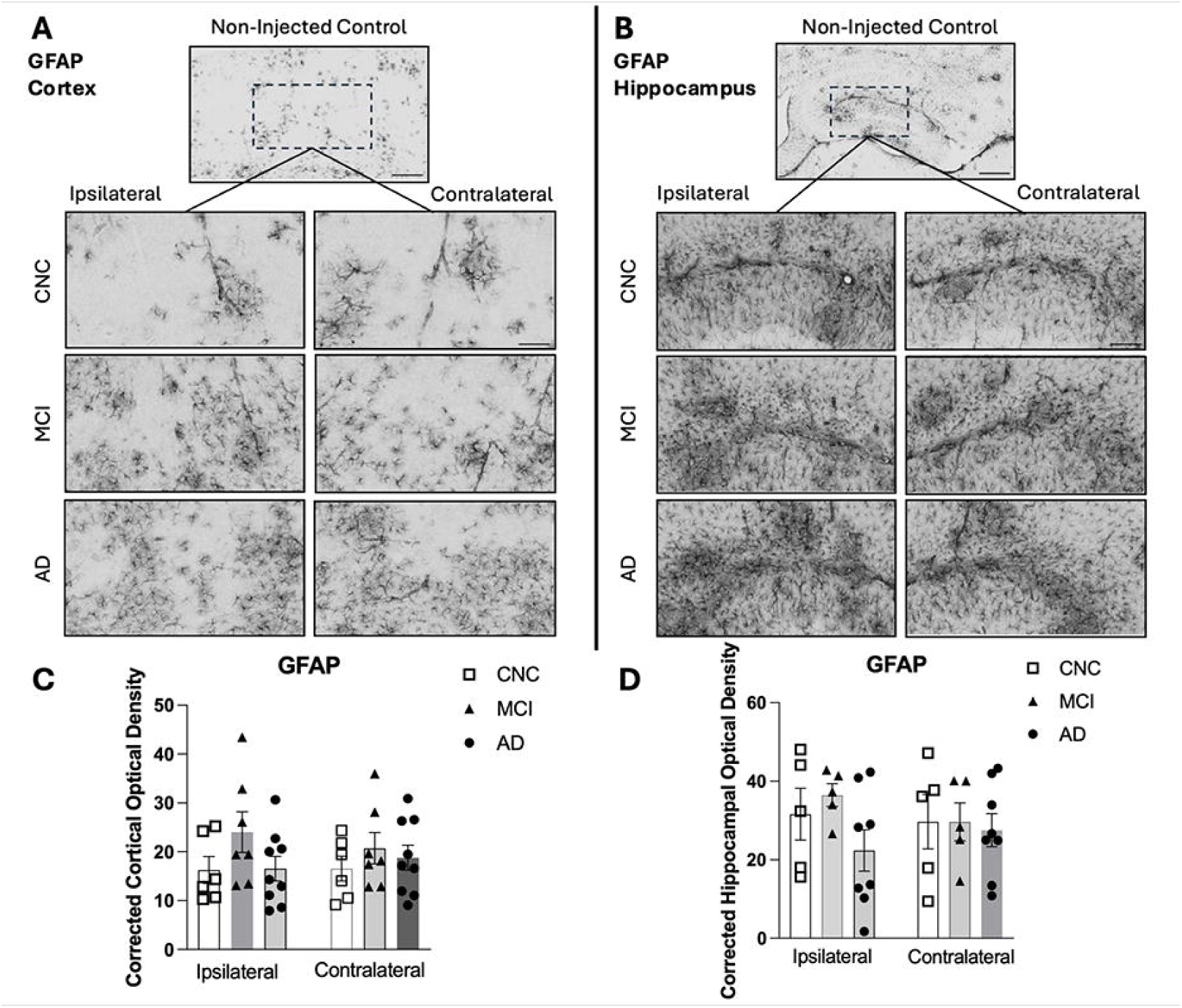
GFAP brightfield immunohistochemical staining of astrocyte-derived extracellular vesicles (AEVs) unilaterally injected into the right hippocampus of female PSAPP mice at 6 months post-injection (mpi). Representative staining of GFAP in the (A) cortex and (B) hippocampus. Scale bars = 250 μm, 150 μm. Depicts the corrected optical densities of 6E10 staining in the (C) cortex and (D) hippocampus. At least 6 mice per group were analyzed. Statistical analyses were performed using One-way ANOVA with a Tukey test. All comparisons were not significant. Abbreviations: CNC, cognitively normal control; MCI, mild cognitive impairment; AD, Alzheimer’s disease; PSAPP, APPswe x PSEN1dE9.

### Biochemical Assessment of Amyloid Burden in the Cortex and Hippocampus of AEV-injected mice

Amyloid expression, detected with antibodies recognizing Aβ/APP N-terminus (6E10), full-length APP (22C11), and Aβ-specific N-terminus (82E1) was quantified via western Blot in the cortex (Figures 6A–6C) and hippocampus (Figures 6E–6G) of AEV-injected mice, normalizing all data to the β-actin loading control. In both the cortex and hippocampus, AD-AEV injected mice showed increased amyloid generation compared with CNC-AEV injected mice. Consistently, 82E1 levels were elevated in the hippocampus of AD-AEV injected mice relative to controls. In contrast, no significant group differences were detected in cortical or hippocampal 22C11 levels across CNC-, MCI-, and AD-AEV injected groups. Statistical analyses confirmed that differences in 6E10, 22C11, and 82E1 expression among groups did not reach significance.

**Figure 6:**
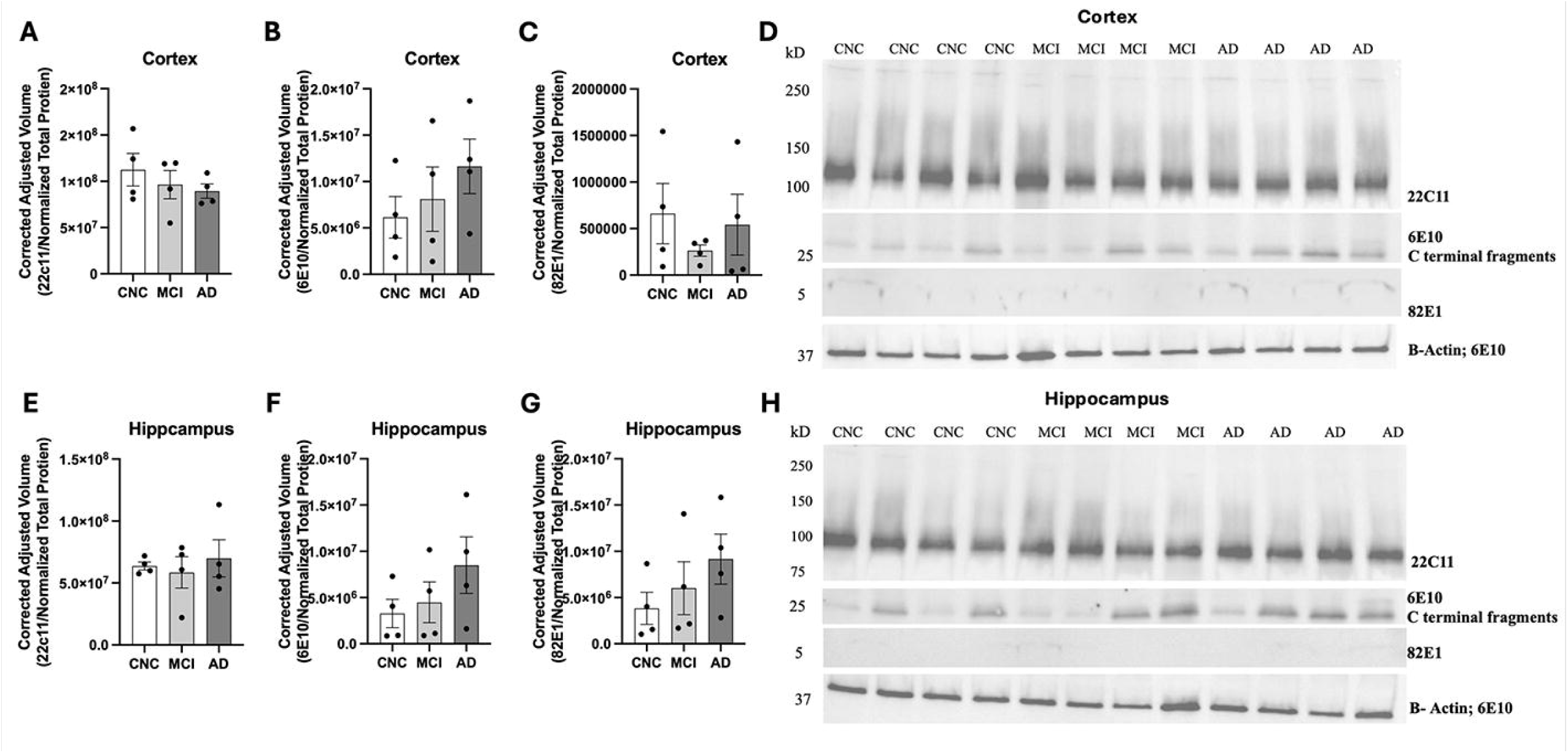
Immunoblots analysis of amyloid presence in the cortex and hippocampus of astrocyte-derived extracellular vesicles (AEVs) unilaterally injected into the right hippocampus of female PSAPP mice. Depicts the corrected, adjusted volume of (A, E) 22C11, (B, F) 6E10, and (C, G) 82E1 protein levels in the (A–C) cortex and (E–G) hippocampus. Representative immunoblots from (D) cortex and (H) hippocampus enriched-regions analyzed with antibodies 22C11, 6E10, 82E1, and β-actin. (n = 3/group, female, 6 months post-injection (mpi)). Statistical analyses were performed using One-way ANOVA with a Tukey test. All comparisons were not significant. Abbreviations: CNC, cognitively normal control; MCI, mild cognitive impairment; AD, Alzheimer’s disease; PSAPP, APPswe x PSEN1dE9.

### Neuropathological and Biochemical Assessment of Tau accumulation in the Cortex and Hippocampus of AEV-injected mice

Previous studies in AD pathology have shown that Aβ can drive tau aggregation and phosphorylation, contributing to neurotoxicity and cognitive decline (Zhang et al., 2021). To de whether AEVs can likewise induce early tau phosphorylation, we assessed phosphorylated tau (p-tau) levels in aged PSAPP mice at 6 mpi using the PHF-1 antibody (Figure 7). Immunostaining revealed no significant increases in p-tau levels in the brains of AEV-injected mice overall. Biochemical quantification by western blot using PHF-1 showed increased cortical and hippocampal p-tau in MCI- and AD-AEV injected mice compared with CNC-AEV injected mice, although these differences did not reach statistical significance.

**Figure 7:**
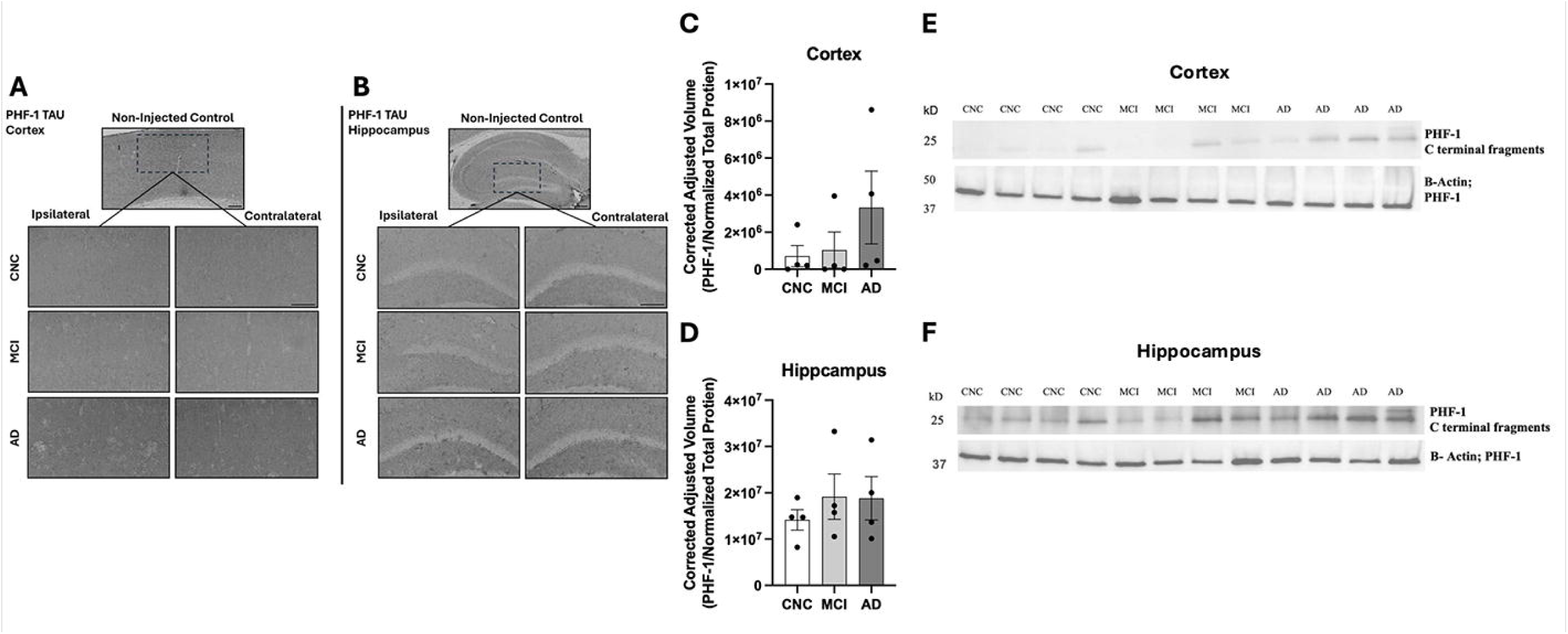
PHF-1 TAU brightfield immunohistochemical staining and immunoblot of astrocyte-derived extracellular vesicles (AEVs) unilaterally injected into the right hippocampus of female PSAPP mice at 6 months post-injection (mpi). Representative staining of PHF-1 TAU in the (A) cortex and (B) hippocampus. Scale bars = 250 μm, 150 μm. Depicts the correct, adjusted volume of (C) PHF-1 TAU in the cortex and (D) hippocampus. Representative immunoblots from (E) cortex and (F) hippocampus enriched-regions analyzed with PHF-1 TAU and β-actin. (n = 3/group, female, 6 months post-injection (mpi)). Statistical analyses were performed using One-way ANOVA with a Tukey test. All comparisons were not significant. Abbreviations: CNC, cognitively normal control; MCI, mild cognitive impairment; AD, Alzheimer’s disease; PSAPP, APPswe x PSEN1dE9.

### Rotarod: Latency to Fall & Fluorescence Immunostaining of Synaptophysin in Cerebellum

Motor coordination, balance, and motor learning were assessed across all AEV-injected groups using the rotarod test. Mice that received MCI- or AD-AEVs exhibited significantly reduced latency to fall compared with CNC-AEV injected mice, indicating impaired motor performance as measured by two-way ANOVA (*p* < 0.0001) (Figure 8A). To explore potential mechanisms underlying these deficits, synaptic density in the cerebellum was evaluated by immunofluorescence using the SY-38 anti-synaptophysin antibody. A reduction in synaptophysin signal was observed in the granule and molecular cell layers of CNC-AEV injected mice, whereas AD-AEV injected mice exhibited higher synaptophysin signal intensity, although these differences did not reach statistical significance (Figure 8B).

**Figure 8:**
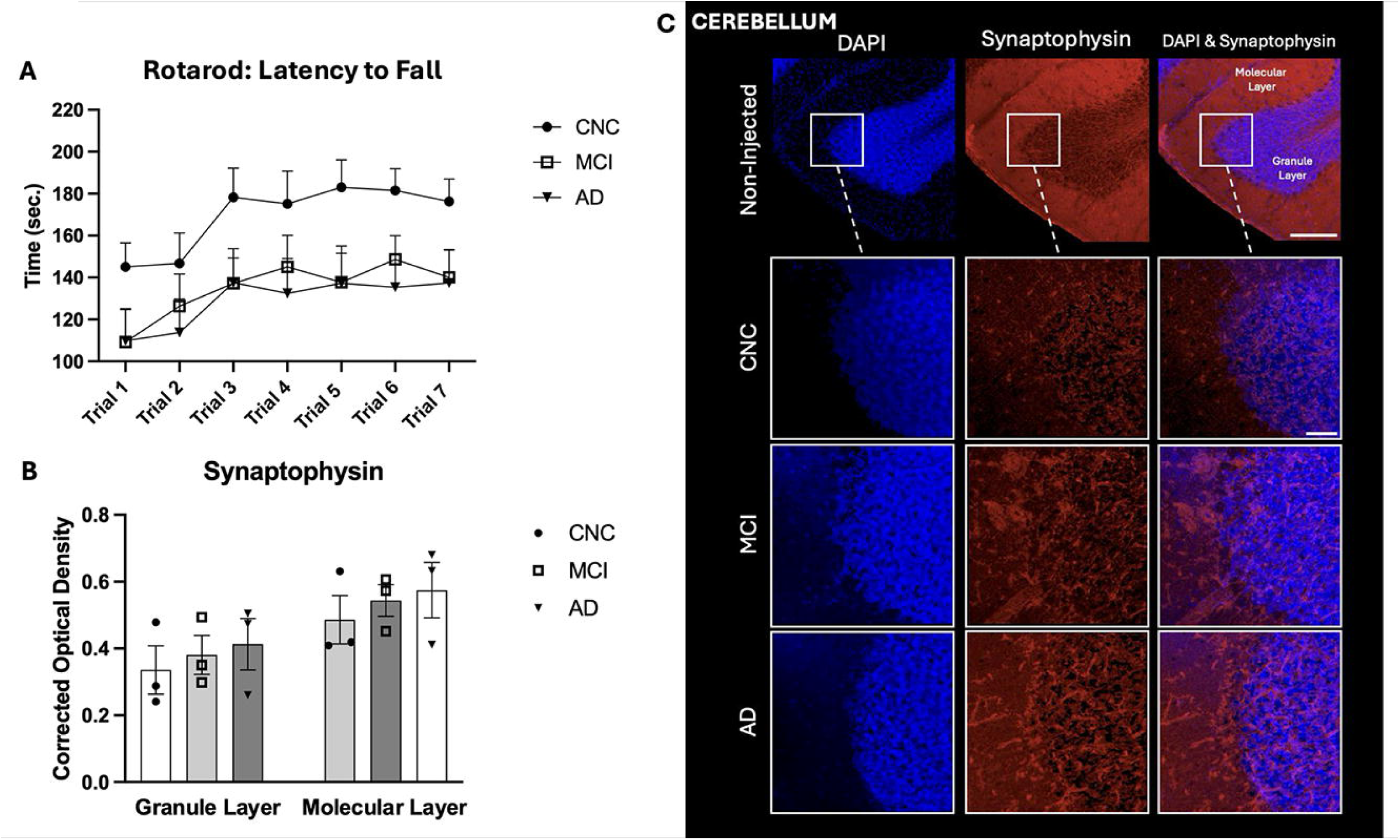
Rotarod behavioral assessment of astrocyte-derived extracellular vesicles (AEVs) unilaterally injected into the right hippocampus of female PSAPP mice and Fluorescence Immunostaining of Synaptophysin in Cerebellum. (A) Depicts the parameter of latency to fall (seconds). Statistical significance was measured by two-way ANOVA statistical test (*p* < 0.0001). (B) Depicts corrected optical density of synaptophysin immunofluorescence staining in the cerebellum. At least three mice per group were analyzed. (C) Representative immunofluorescence staining in the cerebellum using SY-38 antibody. Scale bars = 150 μm, 50 μm. Abbreviations: CNC, cognitively normal control; MCI, mild cognitive impairment; AD, Alzheimer’s disease; PSAPP, APPswe x PSEN1dE9.

## Discussion

Our objective was to investigate the neuropathological and functional impact of AEVs from human plasma in an aged PSAPP mouse model. In the MWM and rotarod assessment, we demonstrated that while plasma AEVs from CNC individuals may possess neuroprotective properties that contribute to memory retention and preserve motor functions, they do not appear to attenuate established amyloid brain pathology or reduce inflammation. Although CNC-AEV injected mice displayed increased motor functions than those injected with MCI- or AD-AEV, synaptic density remained comparable across all groups. Additionally, AEVs from MCI and AD individuals triggered modest early tau phosphorylation, suggesting pathogenic potential analogous to that reported for NEVs. NEVs from individuals with MCI and AD have previously been shown to induce tau accumulation in the brains of C57Bl/6 mice, leading to neuronal degeneration (Winston et al., 2016). Building on this, we sought to expand the investigation from NEVs to AEVs, hypothesizing that AEVs are also implicated in disease progression, as they have been reported to carrying elevated levels of sAPPL and sAPPβ than NEVs (Goetzl et al., 2016). The earlier NEV study used healthy C57BL/6 mice; we instead employed aged PSAPP animals. PSAPP mice are amyloid-dominant yet essentially tau-naïve, developing robust Aβ plaques by nine months. This background allows us to better test whether human AEVs can initiate early tau phosphorylation in the absence of pre-existing tangles, isolating AEV-specific effects and matching the clinical sequence in which extracellular Aβ accumulation precedes tau pathology.

During cued (visible platform) training (Days 1–3; Figure 3), mice injected with MCI- or AD-AEVs showed longer escape latencies and greater path lengths than CNC-AEV controls. Across the hidden (submerged) platform phase (Days 4-7), group performance converged; by Day 7, escape latencies and path lengths were similar across all AEV-injected cohorts (Figure 3A–B). In the Day 8 probe (Trial 1; platform removed), MCI- and AD-AEV groups made fewer entries into the target quadrant and fewer crossings of the former platform location (target platform zone) relative to CNC-AEV mice (Figure 3C & 3E), while time in the target quadrant and total swim distance were comparable (Figure 3D & 3F). Together, these findings indicate stronger spatial memory retention in CNC-AEV injected mice compared with MCI- or AD-AEV injected mice. AEVs have been reported to carry trophic and pro-repair factors in healthy settings (e.g., FGF2, VEGF) that support neurogenesis, survival, and tissue repair, as well as apolipoprotein-D, which has antioxidant and potential geroprotective(Anderson et al., 2018; Dassati et al., 2014; Lange et al., 2016; Li et al., 2015; Pascua-Maestro et al., 2018; Proia et al., 2008)￼ offer an explanation for the relatively better probe performance in CNC-AEV mice. CNC-AEVs may deliver more homeostatic or pro-survival cargo than MCI/AD-derived AEVs, which could carry signals that impair memory consolidation or retrieval. However, this mechanistic link remains inferential in our study and warrants direct cargo profiling and functional validation.

The rotarod test demonstrated that mice receiving AEVs from MCI or AD individuals exhibited significantly shorter latencies to fall than CNC-AEV injected mice, indicating impaired motor coordination and/or motor learning in the MCI- and AD-AEV injected mice (Figure 8A). To examine potential synaptic correlation, we quantified cerebellar presynaptic labeling with SY-38 (anti-synaptophysin) across molecular and granule layers. Contrary to our hypothesis, synaptophysin immunoreactivity did not differ significantly among groups (Figure 8), and the directional ordering, with CNC showing lower labeling than MCI and AD and with AD highest, did not parallel behavior. Structurally, presynaptic terminals in the cerebellar molecular layer predominantly contact Purkinje dendrites (Lindeman et al., 2021; Mavroudis et al., 2019). Prior work shows that synaptic dysfunction emerges early in the AD continuum, including in MCI, often manifesting as decreased synaptic markers (Masliah et al., 2001; Scheff et al., 2007) or electrophysiologic impairments preceding overt plaque deposition in certain models (e.g., Tg2576). (Jacobsen et al., 2006) Conversely, compensatory remodeling can transiently increase synaptic or dendritic metrics despite ongoing pathology (Baazaoui et al., 2017), and spine loss can be accompanied by homeostatic enlargement of remaining spines (Bhembre et al., 2023). While our cohort (9–12-month-old PSAPP mice) and readout (cerebellar presynaptic labeling) differ from those early cortical/hippocampal findings, a compensatory framework could potentially decouple presynaptic abundance from motor performance; effectively, protein levels may remain stable while synaptic efficacy or circuit integration is impaired. The present synaptophysin analysis should therefore be interpreted as preliminary. Notably, the significant motor deficits observed in MCI- and AD-AEV injected mice occurred despite preserved cerebellar synaptophysin density. Synaptophysin staining primarily indexes presynaptic vesicle abundance and serves as a structural correlate rather than a direct measure of synaptic efficacy or plasticity. This dissociation suggests that AEV-induced deficits may be driven by functional circuit-level alterations, which include impaired release probability or disrupted Purkinje cell timing, that are not captured by gross measures of presynaptic protein inventory.

In cortex and hippocampus, immunoblot signals for 82E1 (Aβ N-terminus; β-site-dependent, soluble Aβ peptides) and 22C11 (full-length APP N-terminus) did not differ among groups, whereas 6E10 readouts determined by immunohistochemistry (Figure 4A–B) and immunoblotting (Figure 6B, 6F) were directionally lower in CNC-AEV injected mice than in MCI- or AD-AEV injected mice. Although the 6E10 group differences were not statistically significant, the divergence across antibodies can be informative, leading us to consider that 6E10 (Aβ residues ∼3–8) preferentially reports plaque-associated/aggregated material in tissue, while 22C11 indexes total APP and 82E1 is most sensitive to soluble, β-cleaved Aβ species. Together, these data can suggest that AEV exposure affected deposition or aggregation state rather than overall APP abundance or total soluble Aβ. One interpretation is that CNC-AEVs provide relative protection against plaque accumulation, whereas MCI- and AD-AEVs may carry cargo that facilitates aggregation or stabilizes deposited assemblies. These findings are consistent with reports that inflammatory activation drives astrocytes to release AEVs enriched in Aβ (Li et al., 2020) and that astrocytes carrying Aβ protofibrils secrete neurotoxic Aβ-containing EVs (Söllvander et al., 2016). By contrast, under homeostatic conditions astrocytes have been shown to release EVs bearing trophic factors (e.g., FGF2, VEGF) and apolipoprotein-D, which support neurogenesis, survival, and antioxidative characteristics (Pascua-Maestro et al., 2018; Proia et al., 2008).

In the ipsilateral hemisphere (injection side), MCI-AEV mice showed higher aggregated Aβ plaque burden than AD-AEV mice, although this difference was not statistically significant. In the contralateral hemisphere, plaque burdens were similar between MCI- and AD-AEV injected mice. The hemispheric pattern aligns with findings that amyloid deposition can be asymmetric in preclinical AD and MCI and may become more symmetric as disease progresses to dementia (Kjeldsen et al., 2022; Yoon et al., 2021), which could account for the comparable bilateral burden in the AD-AEV injected mice. Previous studies using bilateral injections of Aβ-containing materials or brain extracts have demonstrated that pathology consistently develops in a focal manner near the infusion site (Langer et al., 2011; Ye et al., 2015). Although our injections were performed unilaterally, the same principle of localized seeding could still be applied. Pathology was more developed in the ipsilateral hemisphere (the injection site) due to direct exposure to the AEV cargo while the contralateral hemisphere served as internal control. Local retention, need-tract-related changes, and immediate immune environment can enhance seeding efficiency and stabilization of deposits near the infusion site (Jean et al., 2015; Ziegler-Waldkirch et al., 2018). By contrast, the contralateral hemisphere (without direct exposure) is expected to show delayed or reduced pathology. Because group differences did not reach significance, and we did not fractionate soluble versus insoluble Aβ or quantify AEV biodistribution, these interpretations are preliminary. Region-matched biochemical assays and in vivo tracking of injected AEVs will be required to determine whether hemispheric differences reflect stage-dependent asymmetry, local AEV effects, or both.

Cortical GFAP immunostaining (Figure 5) was lower in CNC-AEV injected mice than in MCI-and AD-AEV groups, although the group differences did not reach statistical significance. This pattern aligns with the lower cortical Aβ plaque burden (6E10) observed in CNC-AEV mice and is consistent with the possibility that CNC-AEVs carry comparatively less pro-inflammatory cargo (e.g., complement components, cytokines, or miRNAs) or relatively more homeostatic signals, resulting in reduced astrocytic reactivity (Upadhya et al., 2020). We did not perform unbiased proteomic or transcriptomic cargo profiling on the exact AEV pools used for injection; thus, proposed molecular drivers (e.g., complement, cytokine, or miRNA-mediated mechanisms) remain hypotheses that require direct proteomic and transcriptomic characterization in future work. Although a vehicle-only sham cohort was not included, the CNC-AEV group provides a procedure-matched negative control. The negligible gliosis observed in this group, together with prior work from our laboratory (Winston et al., 2016) and others showing only transient needle-track changes after stereotaxic injection ((Song et al., 2013);(McLarnon, 2014)), supports the conclusion that the modest microglial activation seen after AD-AEV delivery reflects vesicle cargo rather than surgical trauma. By contrast, AEVs from MCI and AD donors could bias astrocytes toward a more reactive state associated with greater Aβ deposition. Sampling variability, limited power, or regional heterogeneity remain viable explanations given the absence of statistical significance. In the hippocampus in the ipsilateral hemisphere, GFAP reactivity in CNC-AEV mice was lower than in MCI-AEV mice but higher than in AD-AEV mice. These findings may reflect stage-dependent astrocyte states. MCI-AEVs may acutely enhance GFAP-positive hypertrophy, whereas AD-AEVs could promote a reactive phenotype with reduced GFAP labeling (e.g., dystrophic or GFAP-low states) despite ongoing pathology. Alternatively, high local plaque load or chronic exposure to AD-AEVs may induce cytoskeletal remodeling not captured by GFAP intensity alone. This discrepancy may also reflect the complex role of astrocytes in AD pathology. Unlike amyloid plaque deposition, which progresses in a relatively stage-dependent and regionally predictable manner, astrocytic responses are highly heterogeneous and can vary depending on local microenvironmental cues, cytokine signaling, and regional vulnerability (Kim et al., 2024; Matias et al., 2019). On the contralateral side, GFAP levels were similar across CNC, MCI, and AD groups, an observation not seen in the 6E10 immunostaining. This divergence suggests regional uncoupling between astrocytic reactivity (as measured by GFAP) and plaque metrics. Possible mechanisms include (i) bilateral diffusion of AEVs yielding convergent GFAP responses, (ii) reactive states driven by soluble Aβ or EV cargo independent of plaque density, and (iii) marker limitations as GFAP detects filamentous hypertrophy in only a subset of reactive states, while other markers (e.g., C3, LCN2, vimentin, S100β) may shift without proportional changes in GFAP (Badhwar et al., 2024; Youn et al., 2025). These findings also suggest that astrocytic reactivity in these regions is less dependent on AEV cargo and more influenced by baseline age-related gliosis or PSAPP transgene-driven pathology (Matias et al., 2019). Another possibility is that astrocytic reactivity reaches a saturation point once a threshold of amyloid or inflammatory signaling is crossed, obscuring group-specific differences in GFAP immunostaining (Ceyzériat et al., 2018). Finally, because GFAP primarily marks reactive astrocytes without distinguishing between neuroprotective (A2) and neurotoxic (A1) states, comparable GFAP signals may mask underlying functional differences between astrocyte phenotypes (Liddelow and Barres, 2017). It remains uncertain whether these observations reflect meaningful biological processes or experimental variability. Astrocytes are well established as key regulators of neurodegeneration, capable of exerting both neuroprotective and neurotoxic effects depending on context. Further investigation will be required to determine whether the reduced GFAP signal in CNC-AEV injected mice represents a protective phenotype that facilitates Aβ clearance or, conversely, whether MCI-and AD-AEVs promote a shift toward a reactive, neurotoxic astrocytic state.

PHF-1 immunostaining was absent in cortex and hippocampus across all AEV-injected groups (Figure 7A–B), consistent with the PSAPP model, which develops robust Aβ pathology but lacks overt tau tangle formation. In contrast, PHF-1 immunoblotting revealed higher cortical signals in mice receiving MCI- or AD-AEVs relative to CNC-AEVs (Figure 7C & 7E), although group differences were not statistically significant. Although tau changes were modest and did not reach histological significance, the consistent rotarod impairment suggests a functional consequence that warrants follow-up in models permissive for downstream tangle formation. Because PHF-1 recognizes tau phosphorylated at Ser396/404, the immunoblot increase likely reflects soluble phosphorylated tau species and/or tau fragments rather than mature and/or insoluble aggregates detectable by histology. A potential interpretation is that MCI- and AD-AEVs deliver cargo that enhances tau phosphorylation or stability of phosphorylated species, yielding a biochemical signal without corresponding PHF-1-positive inclusions on tissue sections. This is consistent with previous frameworks in which Aβ-related mechanisms can promote downstream tau phosphorylation in models that do not develop fibrillar tau pathology seen in human AD (Duyckaerts et al., 2008; Héraud et al., 2014; Kitazawa et al., 2012; Seino et al., 2010; Zhang et al., 2021). Taken together, these data suggest that, within the constraints of PSAPP mouse model, tau biochemistry may provide a more sensitive readout of AEV-associated effects than tangle- or plaque-centric histopathology. However, in the absence of detergent-insoluble fractionation, phospho-epitope mapping, or tau seeding assays, mechanistic conclusions remain prefatory. The absence of overt tangles in PSAPP confines our mechanistic claims to early p-tau accumulation rather than mature filament formation. However, recent work validates this approach, demonstrating that APP/PS1-based models are permissive for exogenous tau seeding and readily develop phosphorylated tau after intracerebral delivery of AD-derived pathogenic seeds (Finneran et al., 2024);(Tok et al., 2022)). These studies support our use of an amyloid-rich background as a sensitive platform for detecting the initiating capacity of AEV cargo.

## Limitations

These findings should be interpreted considering several constraints. First, the highly lipophilic nature of extracellular vesicles (EVs) makes them prone to aggregation, preventing uniform delivery across stereotaxic injections, which may have contributed to variability in uptake and effect (Badhwar et al., 2024). This challenge is further compounded by our use of a single unilateral injection, rather than bilateral or systemic delivery. Bilateral paradigms, in which the contralateral hemisphere receives a vehicle, would also have provided an internal sham control to disentangle injection-related effects from AEV biology. Second, we conducted our histological and biochemical analyses with relatively small group sizes, which reduced statistical power and increased variance. Because AEVs were pooled (n = 4) to achieve sufficient protein for dosing, inter-individual variability could not be evaluated. Sample sizes also limited statistical power for biochemical analyses (n = 3 / group), increasing sensitivity to inter-animal variability and reducing our ability to detect modest effects. Accordingly, non-significant group differences are interpreted as exploratory trends rather than definitive negative findings, and conclusions are anchored to statistically significant outcomes.

Additionally, the technical limitations of current EV isolation methods likely introduced heterogeneity. Standard precipitation and immunocapture approaches can co-isolate non-vesicular particles (e.g., protein aggregates, lipoproteins), yielding mixed vesicle populations that complicate our ability to attribute the observed effects solely to AEVs and limit reproducibility across studies (Sigdel et al., 2023). Moreover, we did not perform unbiased cargo profiling on the injected AEV pools. While we employed rigorous immunocapture and quality control to ensure AEV purity, our conclusions regarding specific molecular drivers remain inferential. Future studies employing unbiased proteomic and transcriptomic profiling are necessary to definitively identify the pathogenic cargo species responsibly for the observed phenotypes. We did not image GLAST at single-particle resolution because suitable probes were unavailable; therefore, our interpretations regarding the aggregation state and molecular composition remain provisional. Because EV yield was reserved for in-vivo dosing, we did not repeat TEM and western blot panels on each individual pool. However, higher-throughput capture methods now in development should permit pool-specific ultrastructural confirmation in future studies.

We also lacked a sham (vehicle-only) injection group, preventing complete separation of procedure-related effects from AEV-mediated biology and limits our ability to exclude subtle procedure-related effects. Our choice of animal model imposes additional constraints, as PSAPP mice are primarily amyloidogenic. Our use of an amyloid-dominant, tau-naïve model (PSAPP) means the study captures the earliest tau-seeding events but cannot fully address how AEVs might modulate established tau tangles present in other AD lines. Future work in combined amyloid-and-tau models (e.g., 3xTg-AD or APP/PS1 × P301S crosses) will be needed to verify whether the vesicle effects observed here extend to brains with pre-existing tauopathy (Prikhodko et al., 2020) (JAX #004462). Consequently, extrapolation to the whole human AD spectrum should be cautious. Finally, assay-specific considerations apply. The 6E10 antibody recognizes an Aβ epitope present on multiple APP-derived species (Baghallab et al., 2018), whereas 22C11 (full-length APP) and 82E1 (β-site-dependent Aβ) assess distinct pools of APP (Hunter and Brayne, 2017; Readel et al., 2023). Without biochemical separation of soluble versus insoluble fractions and use of conformation-specific reagents, amyloid “burden” cannot be entirely attributed to plaques. Likewise, synaptophysin provides only a proxy for presynaptic abundance and does not measure synaptic function (e.g., efficacy, plasticity, or circuit timing); therefore, we cannot exclude functional synaptic or circuit-level deficits as contributors to the observed motor impairment. PHF-1 immunoblots detect phospho-tau epitopes that may not directly correspond to neurofibrillary tangles detectable by histopathology (Limorenko and Lashuel, 2022).

## Future directions

Future research should build on these findings by addressing the outlined constraints and exploring AEV biology in greater mechanistic depth. Comprehensive cargo profiling, including proteomic, transcriptomic, and conformation-specific assays, will be crucial in defining the molecular drivers of AEV effects. Although our measures meet current MISEV2018 purity standards, future work will incorporate gradient or SEC purification to provide an orthogonal confirmation of vesicle-associated GLAST. Future studies should employ labeled anti-GLAST antibodies or ExoView platforms as they become accessible. In particular, astrocytes can transition to reactive A1 phenotypes under AD conditions, releasing AEVs enriched in complement proteins (Goetzl et al., 2018). Complement proteins (e.g. C1q, C3) have been implicated in synaptic pruning and early synapse loss in multiple mouse and human studies of Alzheimer’s disease (Hong et al., 2016; Shi et al., 2015) and may mediate microglial or astrocytic engulfment of synapses. In parallel, AEVs can carry regulatory miRNAs such as miR-155-5p (shown in SOD1^G93A^ ALS models) that reduce neuronal survival and neurite growth, illustrating how non-protein cargoes may contribute to neurotoxicity (Marton et al., 2023). This highlights their potential to reshape neuronal pathways through non-protein cargo. By defining the relative contributions of these molecular classes, we can clarify whether AEVs are predominantly neuroprotective or neurotoxic in various contexts. To address kit-related caveats (co-precipitation of lipoproteins), we plan to incorporate size-exclusion chromatography followed by GLAST capture, in line with recent comparisons showing higher purity and yield (Figueroa-Hall et al., 2025).

Methodologically, future studies should incorporate sham-injected controls, including bilateral injections where one hemisphere receives AEVs, and the contralateral hemisphere receives a vehicle as an internal control. However, unilateral intrahippocampal injection is a reductionist exposure model and does not represent physiological EV biodistribution. Because EVs may traffic beyond the injection site, the contralateral hemisphere cannot be assumed to be fully EV-naive without direct biodistribution measurements; therefore, hemispheric comparisons are interpreted as reflecting relative, not absolute, exposure. Systemic or repeated injections should also be considered to better approximate physiological EVs distribution and to evaluate whether the observed effects extend beyond a single site. Future studies will include vehicle, CNC-AEV, and AD-AEV groups to fully disentangle cargo-specific outcomes from surgical influences. Future studies will increase cohort sizes for biochemical endpoints and include independent biological replication (e.g., multiple independently prepared donor pools per group) to improve power and generalizability. Region-matched biochemical assays and soluble/insoluble fractionation will further reduce variability and strengthen interpretation. We acknowledge that the model cannot address later propagation across a tau-rich network; future studies will therefore extend these findings in combined amyloid- and tau-expressing lines. Utilizing complementary animal models that recapitulate both amyloid and tau pathology will be essential to observe the full scope of AEV-mediated effects. Mutant mouse lines (e.g., 3xTg-AD mice that express both humanized APP (Swedish) and tau mutations (PSEN1 (M146V), and tau (P301L)) may provide a more physiologically relevant context for assessing whether AEVs exacerbate the synergistic interaction between amyloid deposition and tau aggregation observed in human AD (Jiang et al., 2024; Tag et al., 2022). Examining AEVs within 3R/4Rtau-mutant bigenic mice could also be another possibility to clarify their impact in propagating different forms of tau (Spencer et al., 2025). In parallel, tau knockout (MAPT-/-) mice would serve as a negative control to confirm whether the tau pathology we observed is directly induced by AEV exposure rather than secondary to endogenous tau dynamics (Denk and Wade-Martins, 2009). Demonstrating that AEV-treated tau knockout mice fail to develop tau pathology would potentially implicate AEV cargo as the pathogenic driver of tau-related alterations. Additionally, incorporating C57BL/6 wild-type and tau knockout mice as comparators to our PSAPP cohort would more clearly define the specific contribution of Aβ pathology to AEV-mediated tau seeding and propagation. Extended post-injection survival or combined amyloid- and tau-expressing lines will also be necessary to establish whether AEV-induced phosphorylation evolves into overt tau aggregation. Together, these complementary models would provide more comprehensive platforms and contextual understanding of whether AEV exposure amplifies existing tauopathy, triggers tau pathology de novo, or acts primarily through amyloid-dependent mechanisms. Integrating these systems would therefore advance mechanistic understanding of how human-derived AEVs contribute to the amyloid-tau axis central to AD progression.

Finally, electrophysiological analysis of neuronal function (Chun et al., 2021; Kuo et al., 2014) and ultrastructural mapping of synapses (Reilly et al., 2017) will help establish whether AEVs primarily alter synaptic abundance, efficacy, or circuit-level properties. Future advances in magnetic-net and other high-yield capture methods (Suresh and Zhang, 2025); Rarokar et al., 2024) are also expected to yield enough material from sub-milliliter plasma volumes to support single-donor studies, an approach we aim to adopt in follow-up experiments. Together, these efforts will advance understanding of how AEVs influence the course of AD and may inform their development as biomarkers or therapeutic targets.

## Conclusion

Our study demonstrates that AEVs from individuals with MCI and AD exacerbate motor and cognitive impairments in aged PSAPP mice, while AEVs from cognitively normal individuals may provide partial neuroprotection. Although CNC-AEVs were associated with improved behavioral outcomes, they did not reduce established Aβ pathology. Plasma MCI- and AD-AEVs induced modest early tau phosphorylation in an amyloidogenic mouse model of AD (PSAPP). These findings highlight the dual and context-dependent nature of AEVs, underscoring their potential role as both propagators of neuropathology and carriers of neuroprotective cargo. By extending prior work on NEVs, this study establishes AEVs as active participants in disease-related processes, including tau propagation and motor dysfunction. Future work incorporating larger cohorts, sham and systemic injection paradigms, and comprehensive cargo profiling will be essential to define the molecular determinants of these divergent effects. Ultimately, a deeper understanding of AEV biology may enable the development of EV-based biomarkers and therapeutic strategies aimed at modifying the course of AD.

## Permission to reuse and Copyright

Permission must be obtained for use of copyrighted material from other sources (including the web). Please note that it is compulsory to follow figure instructions.

## 2 CRediT authorships contribution statement

**Bao Quach**: Formal Analysis, Investigation, Writing - original draft, Writing - review & editing. **Sahar Salehi**: Formal Analysis, Investigation, Writing - review & editing. **Rejina Roufegarinejad**: Formal Analysis, Investigation, Writing - original draft. **Michael Mante**: Investigation. **Jazmin Florio**: Investigation. **Robert A. Rissman**: Conceptualization, Formal Analysis, Methodology, Investigation, Projection Administration, Writing - review & editing. **Charisse N. Winston**: Conceptualization, Formal Analysis, Methodology, Investigation, Projection Administration, Writing - review & editing.

## 4 Declaration of Generative AI and AI-assisted technologies in the manuscript preparation process

During the preparation of this work, the author(s) used ChatGPT in order to enhance readability and review the grammar of some sections. After using this tool/service, the author(s) reviewed and edited the content as needed and take(s) full responsibility for the content of the published article.

## 5 Acknowledgements

The authors gratefully acknowledge the UC San Diego Shiley-Marcos Alzheimer’s Disease Research Center (ADRC) for providing the human plasma samples used in this study. We also thank the Epstein Family Alzheimer’s Therapeutic Research Institute at the University of Southern California for providing the essential resources and infrastructure required to conduct this research.

## 6 Data Availability Statement

The original contributions presented in the study are included in the article/supplementary material; further inquiries can be directed to the corresponding author/s.

